# Lifecourse sex-specific molecular response to early-life exposures of toxic substances

**DOI:** 10.64898/2026.08.20.746014

**Authors:** Shuhua Fu, Benpeng Miao, Wanqing Shao, Cristian Coarfa, Ravindra Kumar, Prashant Kumar Kuntala, Bongsoo Park, Sandra l. Grimm, Rahul Jangid, Justin A. Colacino, Xiaoyun Xing, Daofeng Li, Shaopeng Liu, Robert B. Hamanaka, Claudia Lalancette, Maureen A. Sartor, Christopher Krapp, Gregory E Crawford, Heather B. Patisaul, Tim Wiltshire, TaRGET II Consortium, David Aylor, Shyam Biswal, Gökhan M. Mutlu, Sanjay Rajagopalan, Wan-Yee Tang, Dana C. Dolinoy, Ting Wang, Marisa S. Bartolomei, Cheryl L. Walker, Laurie K. Svoboda, Bo A. Zhang

## Abstract

Toxicants in the environment can significantly impact physiology. Environmental chemical exposures during early developmental stages disturb normal embryonic development and programming, and dramatically impact long-term health as individuals age. Female and male animals show distinct phenotypes when responding to a given chemical exposure. Here, through the TaRGET II (Toxicant Exposures and Responses by Genomic and Epigenomic Regulators of Transcription) consortium, we systematically explored sex-specific transcriptomic and epigenomic alterations in response to various toxicants, including arsenic (As), lead (Pb), tributyltin (TBT), bisphenol A (BPA), di(2-ethylhexyl) phthalate (DEHP), dioxin (TCDD), and fine particulate matter (PM2.5), across three time points in mice exposed two weeks prior to conception through gestation and lactation. After being exposed to toxicants during the embryonic and early postnatal developmental stages, 1,025 omics datasets were generated from the liver and analyzed across three mouse life stages. We discovered a significant sex-biased molecular response to distinct exposures in the liver at both the transcriptomic and epigenetic levels, showing dynamic changes across mouse development and aging. The perturbed pathways and transcription factors in response to different chemical exposures in both sexes were further evaluated to measure the sex-specific impact of each toxic exposure in the liver. Overall, this study presents the most detailed investigation of sex-specific molecular signatures under the influence of developmental exposures to toxic substances.

## Introduction

Mammals exhibit pronounced sex differences in development, physiology, and behavior, which are further distinguished at the molecular level, including metabolism, immune responses, neurological function, and disease susceptibility^1–6^. These sex-specific traits are primarily driven by differences in sex chromosome composition (XX vs. XY) and the action of sex hormones, particularly androgens and estrogens^7–9^. These hormones regulate gene expression directly and shape the epigenetic landscape during critical developmental windows, such as early postnatal life and puberty^10,11^. Over time, these epigenetic regulatory mechanisms establish and maintain sex-specific gene expression patterns across various tissues^12,13^. Particularly, the liver undergoes pronounced sex-specific morphogenesis during postnatal development, with sex-biased gene expression patterns that directly shape sex-specific physiological functions, including xenobiotic metabolism, lipid and steroid processing, immune responses, and hormone regulation^14–16^. Liver sex morphogenesis is believed to occur primarily between postnatal weeks 2 and 8, driven by differences in sex hormone levels (testosterone in males, estrogen in females) and sex-specific growth hormone secretion patterns^17^. These hormones act through their respective receptors, i.e., testosterone activates the androgen receptor (AR), while estrogen primarily binds to estrogen receptor alpha (ERα), inducing differential transcriptional programs^12–16^. For instance, the male liver expresses genes such as *Cyp3a11* and *Cyp2c29*^18,19^, which are involved in steroid and amino acid metabolism, whereas the female liver preferentially expresses *Cyp2b9* and *Cyp7a1*, which are important for lipid metabolism and bile acid synthesis^20,21^.

As a central organ for metabolism and detoxification, the liver is a primary target of damage following toxicant exposure. Due to inherent molecular and hormonal differences between males and females, the liver exhibits sex-specific responses and susceptibilities to toxicant-induced injury^22,23^. These differences are largely driven by sex-specific regulation of key hepatic functions, including xenobiotic metabolism, inflammatory signaling, and oxidative stress responses^14–16,22,23^. In particular, early-life exposure to environmental toxicants, such as endocrine-disrupting chemicals like bisphenol A (BPA), 2,3,7,8-tetrachlorodibenzo-p-dioxin (TCDD), and di(2-ethylhexyl) phthalate (DEHP); metals or metalloid (e.g., arsenic, lead); and airborne particulates (e.g., PM_2.5_), can interfere with sex-specific liver development, or sex morphogenesis, resulting in long-lasting sex-biased dysregulation and increased disease susceptibility in a sex-dependent manner^24–31^. For example, developmental exposure to TBT has been shown to cause fatty liver disease in male mice^32^. Maternal BPA exposure induced sex-specific and dose-specific changes in the expression of genes regulating inflammation and mitochondrial function of the pancreatic islets of F1 and F2 mice offspring^33^. TCDD and arsenic (As) exposure disrupted male-biased lipid and steroid metabolism, increasing the risk of nonalcoholic fatty liver disease^34–36^. BPA exposure induced female-specific overexpression of *Cyp2b9*, and further altered lipid homeostasis, potentially leading to steatosis^25,37^. Recent studies indicated a stronger respiratory-related health outcome in females due to exposure to air pollution^38–40^. Exposure to traffic-related air pollution and chronic obstructive pulmonary disease (COPD) resulted in higher chances of inflammatory disease of the respiratory airways among females^41^.

Toxicant exposure can cause both acute damage and long-term chronic effects in animals, often in a dose-dependent manner^42–44^. Even at low environmental levels, toxicants have been shown to induce reproductive impairments, carcinogenic outcomes, and organ-specific dysfunction, including damage to the liver, brain, and reproductive organs^27,29–33,45–53^. Importantly, females and males are affected differently, largely due to sex-specific hormone production, differences in metabolism, and immune responses, leading to differential sensitivity to environmental toxicants. Although the sex-specific impacts of various toxicants have been studied in diverse experimental setups and animal models, most research has focused on single-exposure scenarios. As a result, it remains largely unclear how distinct toxicants uniquely disrupt gene regulation and the epigenetic landscape in a toxicant-specific and sex-specific manner. Furthermore, how early-life exposures affect male and female animals differently across developmental stages is still poorly understood, highlighting the need for comprehensive, multi-exposure, and sex-aware toxicological studies.

In this study, we systematically analyzed a comprehensive multi-omics dataset generated by the TaRGET II consortium^54^, which profiles the molecular effects of early-life exposure to diverse environmental toxicants in mice. Specifically, we examined the liver’s response to seven well-characterized toxicants: arsenic (As), lead (Pb), bisphenol A (BPA), tributyltin (TBT), di(2-ethylhexyl) phthalate (DEHP), tetrachlorodibenzo-p-dioxin (TCDD), and fine particulate matter (PM2.5) air pollution. The sex-balanced experimental design of this dataset enabled us to systematically investigate sex-specific patterns of gene expression, chromatin accessibility, and DNA methylation across various exposures. Furthermore, we analyzed how these molecular signatures vary across three key life stages, including early postnatal (3 weeks), young adult (5 months), and later adult (10 months), to capture the long-term and dynamic effects of early-life toxicant exposure. Finally, we evaluated how these exposures disrupt normal sex-specific gene regulatory programs in the liver, revealing the extent to which environmental toxicants may reshape the molecular architecture of this vital metabolic organ in a sex- and exposure-specific manner.

## Result

### Comprehensive molecular changes in response to early-life environmental toxicant exposure in female and male mice

TaRGET II consortium was designed to systematically investigate the transcriptional and epigenetic alterations in response to early-life exposures to various environmental toxicants in both female and male animals at different life stages across multiple tissues. Environmental toxicants, including As, Pb, BPA (high and low dose), TBT, DEHP, TCDD, and PM_2.5_ air pollutions (two exposures with distinct compositions generated by John Hopkins University (JHU) and University of Chicago (CHI)), were independently maternally exposed to the embryo from 2 weeks before pregnancy until postnatal 3 weeks, then the exposed animals and matched non-exposed controls were maintained without toxicant exposure until 10 months old (**Fig. 1a, methods**). To detect the molecular changes in response to distinct environmental toxicants in both females and males, we analyzed 493 RNA-seq, 346 ATAC-seq, and 186 whole genome bisulfite sequencing WGBS data from liver samples of weaning (3 weeks), early adulthood (5 months), and late adulthood (10 months). For each exposure, multiple replicates from both female and male mice were used to detect high-confidence molecular changes (**Supplementary table 1**).

**Figure 1.**
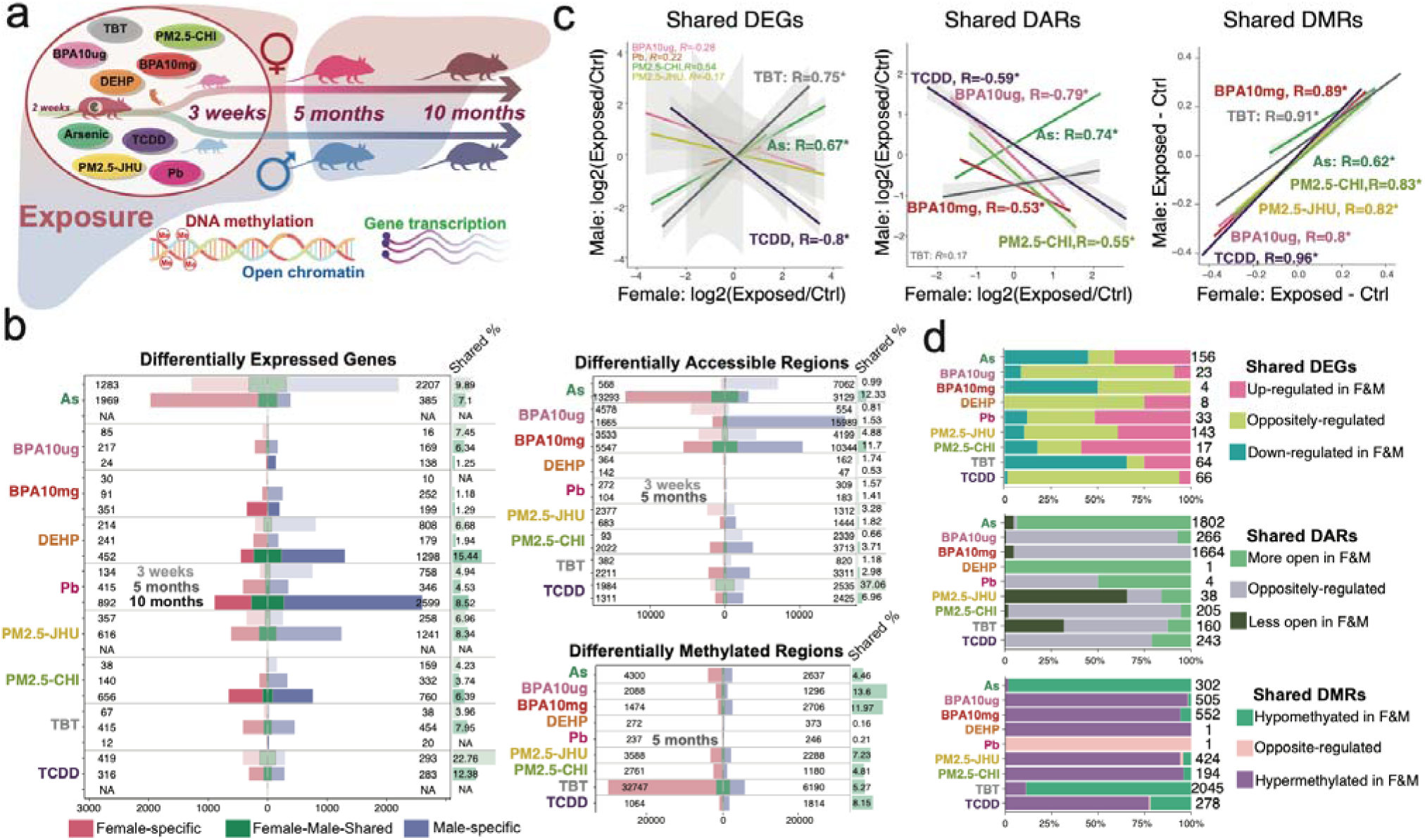
Early-life environmental exposures induce molecular changes in both female (F) and male (M). **a)** Schematic overview of the study workflow. The exposure window was from 2 weeks pre-conception through pregnancy until weaning at 3 weeks of age. **b)** Numbers of female-specific(pink), male-specific(blue), and female-male-shared(green) differentially expressed genes (DEGs at 3 weeks, 5, and 10 months), differentially accessible regions (DARs at 3 weeks and 5 months), and differentially methylated regions (DMRs at 5 months) in liver across different exposures. Green bars on the right side indicate the percentage of female-male-shared molecular features in response to a single exposure. **c)** Correlation of signal changes of female-male-shared DEGs, DARs, and DMRs in female (x-axis) and male (y-axis). **d)** The distribution of changing concordance of female-male-shared DEGs, DARs, and DMRs.

We first identified differentially expressed genes (DEGs), differentially accessible regions (DARs), and differentially methylated regions (DMRs) induced by distinct toxicant exposures in liver, compared to their age-matched normal controls (**Fig. 1b, Supplemental tables S2-S7**). To better characterize the sex-specific differences in molecular changes induced by toxicant exposures, the molecular changes of sex chromosomes (chrX and chrY) were left out, and only changes on autosomes were kept for downstream analysis. At each life stage, strong sex-specific molecular responses to toxicant exposure were observed. At the weanling stage, exposures to As, DEHP, Pb, and PM_2.5_-CHI induced significantly more gene expression changes (DEGs) in males, whereas PM_2.5_-JHU and TCDD exposures induced more DEGs in females. At 5 months of age, exposures to BPA10mg and both PM_2.5_ exposures resulted in more DEGs in males, with only As exposure inducing significantly more DEGs in females. In 10-month-old mice, DEHP and Pb exposures induced significantly more gene changes (DEGs) in males. The genes that changed expression in both sexes under the same toxicant exposure accounted for a small fraction compared to the sex-specific DEGs in response to the exposure (**Fig. 1b**). As exposure induced the most female-male-shared DEGs at 3 weeks and 5 months, whereas DEHP and Pb exposures induced the most female-male-shared DEGs at 10 months old.

The exposure-induced epigenetic alterations revealed pronounced sex-specific patterns, with limited overlap in signatures between females and males (**Fig. 1b**). Compared to other environmental toxicants, DEHP and Pb exposures resulted in relatively modest changes in chromatin accessibility and DNA methylation. At 3 weeks of age, exposures to As, BPA, PM_2.5_-CHI, and TBT induced detectable changes in chromatin accessibility. At 5 months of age, these changes became more extensive, indicating long-term epigenetic remodeling that propagated well beyond the exposure window. BPA exposure led to a notably higher number of male-specific DARs, whereas arsenic exposure induced more female-specific DARs, highlighting sex-dependent epigenetic responses. TBT exposure, in particular, resulted in substantial DNA methylation changes, especially in females. In the female liver, a total of 32,747 differentially methylated regions (DMRs) were identified, with ∼95% showing hypomethylation. These findings emphasize the marked sex-specific nature of epigenetic responses to environmental exposures and underscore the potential for long-term consequences on gene regulation and liver function.

Given the limited number of molecular signatures shared between females and males in response to the same exposure, we further analyzed the concordance of these molecular signatures in both sexes (**Fig-1c, S-Fig-1a**). Our results first revealed that gene expression changes in males and females were largely concordant at 3 weeks of age, reflecting a shared molecular response to acute toxicant exposure (**S-Fig 1a**). At 5 months old, gene expression changes in males and females were less concordant, only As, TBT, and TCDD-induced shared differentially expressed genes (DEGs) exhibited significant correlation between the two sexes at 5 months (**Fig. 1c, S-Fig. 1 b)**. In contrast, epigenetic changes, particularly DNA methylation modifications, demonstrated a strong and significant correlation between males and females. Except for Pb and DEHP, all other exposures induced female-male-shared DNA methylation changes, which showed strong positive correlations between the sexes (**Fig-1c)**. Furthermore, we observed consistent correlations for TCDD-induced shared DEGs and DARs, both showing negative correlations, indicating that the same genes and regulatory elements were oppositely regulated in a sex-specific manner at 5 months (**Fig. 1d**). Interestingly, TCDD-induced shared DEGs and DARs at 3 weeks old were positively concordant (**S-Fig. 1c**). As-exposure-induced shared molecular signatures revealed a significantly strong positive correlation and concordance between males and females for both epigenetic and transcriptional changes, especially 95% of DARs were more open and 99% of DMRs were hypermethylated in both sexes at 5 months (**Fig. 1d**), suggesting As-exposure impacted the liver with a potentially unisex regulatory mechanism. Other toxicants, particularly BPA and PM2.5, induced strongly concordant hypermethylation changes in female–male shared DMRs (**Fig. 1d**). These results suggested that DNA methylation changes can be used as biomarkers to detect early developmental exposure to environmental toxicants in mice.

### Early-Life Exposure Induces Pathway-Level Perturbations in the Mouse Liver

Given the pronounced sex-specific molecular changes, we performed KEGG pathway and Gene Ontology (GO) enrichment analyses to better understand the dysregulated pathways and biological processes in the mouse liver following early-life exposure to environmental toxicants. Enrichment analyses were conducted separately for differentially expressed genes in females and males across three life stages. The results revealed persistent sex-specific pathway perturbations at all time points (**Fig 2a, S-table S8-10**). Notably, male animals exhibited a greater number of dysregulated pathways compared to females. As-exposure induced significant pathway alterations in 3-week-old males and 5-month-old females. Both DEHP and Pb exposures caused substantial pathway dysregulation in male liver, particularly at 3 weeks and again at 10 months of age. In contrast, PM_2.5_-JHU and TBT exposures led to more pathway disruptions primarily in 5-month-old males.

**Figure 2.**
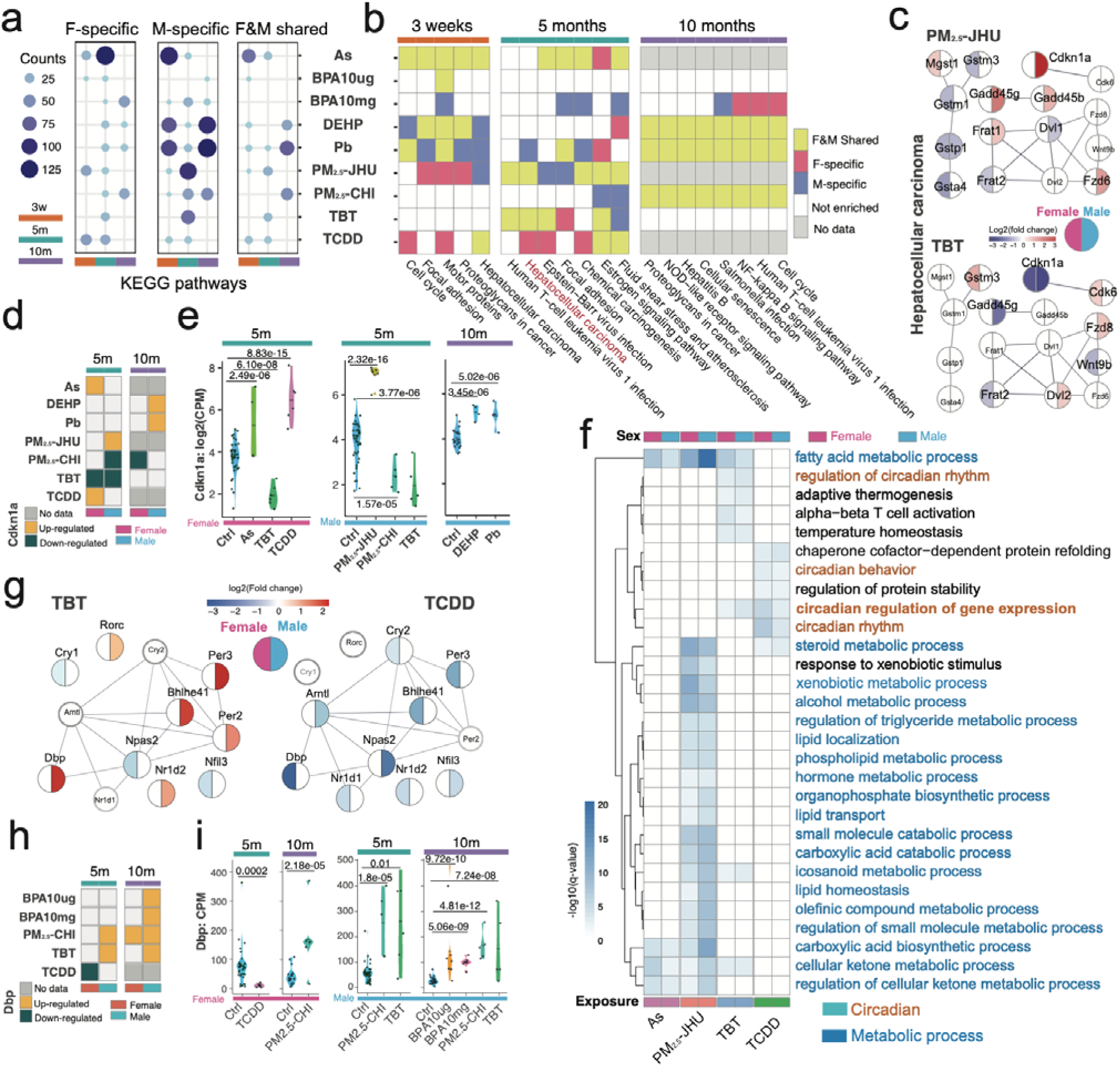
The perturbed KEGG pathways in female and male liver by different exposures. **a)** The number of female-specific (left), male-specific (middle), and female-male-shared (right) KEGG pathways that were perturbed by early-life toxicant exposures at three life stages. **b)** The heatmap of female-male-shared KEGG pathways that were commonly perturbed by at least two early-life toxicant exposures at three life stages. **c)** The protein-protein-interaction network of genes that are dysregulated in the hepatocellular carcinoma pathway in 5-month-old females and males under _PM2.5_-JHU and TBT exposure. **d**) Expression changing pattern and **e)** expression plot of Cdkn1a in 5-month-old and 10-month-old liver across exposure conditions. The p-values from the binomial test were shown for significance. **f)** The heatmap of female-male-shared GO biological processes that were commonly perturbed by at least two early-life toxicant exposures at three life stages. **g)** The protein-protein-interaction network of genes that are dysregulated in the circadian regulation of gene expression in 5-month-old females and males under TBT and TCDD exposure. **h**) Expression changing pattern and **i)** expression plot of Dbp in 5-month-old and 10-month-old liver across exposure conditions.

In addition to the strong sex-specific dysregulation of KEGG pathways, certain pathways were perturbed in both females and males, and even across multiple exposures (**Fig. 2b**). At 10 months of age, DEHP, Pb, and PM_2.5_-CHI exposures induced similar alterations in pathways related to cell cycle regulation, cellular senescence, immune response, and infection, in both sexes. While earlier developmental stages exhibited more exposure-specific pathway changes, some pathways were frequently affected regardless of exposure, including focal adhesion and hepatocellular carcinoma pathways (**Fig. 2b**). At 5 months of age, the hepatocellular carcinoma pathway was notably impacted by both PM_2.5_-JHU and TBT exposures in males and females. To further explore this, we examined the expression patterns of genes within the hepatocellular carcinoma pathway (**Fig 2c**). Interestingly, only a limited number of genes showed concordant changes across sexes and exposures, such as *Gstp1* (PM_2.5_-CHI) and *Cdkn1a* (TBT). The majority of genes, however, displayed strong sex-specific dysregulation, despite being part of commonly altered pathways. These findings suggest that while certain biological pathways are consistently affected across sexes and exposures, sex-specific transcriptional responses play a dominant role in shaping the molecular outcome of early-life toxicant exposure. We also examined the gene expression changes of *Cdkn1a* (**Fig. 2d, e**), which encodes the cyclin-dependent kinase inhibitor p21, a protein with a multifaceted role in liver function, particularly in the context of metabolic dysfunction-associated steatotic liver disease (MASLD)^55^. *Cdkn1a* expression is known to be influenced by various environmental exposures^56,57^. Following TBT exposure, p21 expression was significantly downregulated in both females and males at 5 months of age but remained unchanged at later stages. Other exposures, however, induced changes in p21 expression in a sex-specific manner, suggesting that each environmental toxicant may uniquely modulate the regulation of this key gene.

Gene Ontology (GO) analysis of exposure-induced differentially expressed genes (DEGs) (**S-table S11-13, Fig S2-3**) revealed similar sex-specific patterns as those observed in dysregulated KEGG pathways. A strong sex-specific pattern was evident, and only a limited number of biological processes were shared by both females and males across multiple exposures (**Fig 2f, S-table S14**). Most of these shared biological processes pertained to metabolic processes and circadian rhythm regulation. PM_2.5_-JHU exposure induced a significant number of dysregulated metabolic processes in both females and males. Fatty acid metabolic process and cellular ketone metabolic process were enriched in both sexes under exposure to As, PM_2.5_-JHU, and TBT (**Fig 2f**). Meanwhile, although TBT and TCDD dysregulated circadian rhythm pathways in both sexes, gene expression associated with circadian rhythm still exhibited a sex-specific pattern (**Fig 2g**). D-box binding protein (*Dbp*) plays a crucial role in circadian rhythm regulation in the liver, and its expression oscillates with a daily rhythm^58,59^. We observed that *Dbp* expression was significantly upregulated in males by multiple toxicant exposures, but showed minor changes in females (**Fig. 2h**). Notably, PM_2.5_-CHI and TBT exposure dramatically upregulated *Dbp* expression in both 5-month-old and 10-month-old males (**Fig 2i**). However, in females, TCDD seemed to downregulate the expression of *Dbp* and other genes such as *Per3* and *Bhlhe41*, which were unaffected in males; another set of genes (*Arntl*, *Npas2*, *Nfil3*) were also downregulated (**Fig 2g**).

### Sex-specific enrichment of transcription factor binding motifs in the differential epigenetic regions

Epigenetic modifications are tightly associated with gene regulatory elements and serve as a bridge connecting gene regulation with environmental stimuli, often mediated through transcription factor (TF)-genome interactions. To better understand sex-specific epigenetic changes in response to environmental stimuli (**Fig 1b**), we performed *de novo* motif analysis for differentially accessible and methylated regions in females and males separately. Interestingly, we observed strong sex-specific enrichment of TFs in both more open and closed differentially accessible regions (DARs) induced by each exposure, with very few TFs shared between males and females (**Fig 3a; S-Fig-4, a-d; S-table S15-16**).

**Figure 3.**
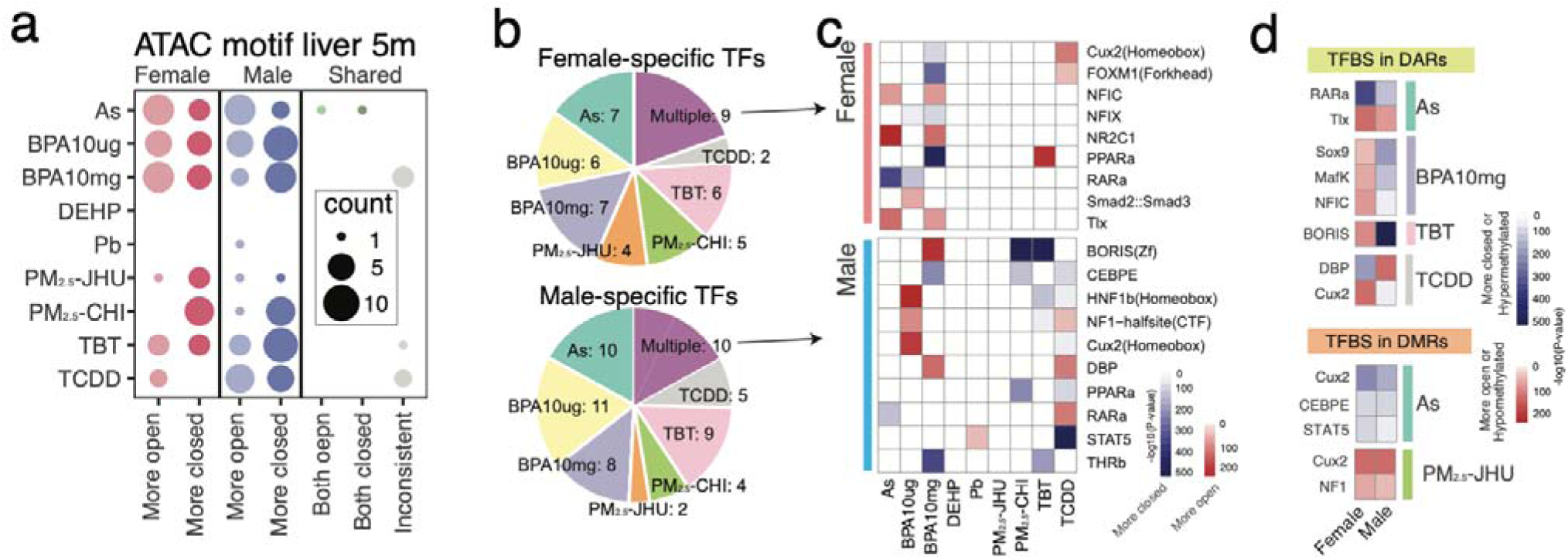
Enriched TF binding motifs in DARs of female and male liver. **a)** The number of female-specific (left), male-specific (middle), and female-male-shared (right) TF binding motifs that are enriched in DARs in response to early-life toxicant exposures at 5 months of age. **b)** Distribution of female-specific (top) and male-specific (bottom) TF binding motifs across distinct exposures. **c)** Heatmap of enrichment significance (p-value) of common TF binding motifs enriched in DARs of at least two exposures. **d)** Female-male-shared TF binding motifs enriched in DARs and DMRs in female and male liver.

Detailed exploration of sex-specific TFs indicated that the enriched TFs were strongly exposre-specific, with only 9 or 10 TFs commonly appearing across multiple exposures (**Fig 3b**). For instance, *Nfic*, *Nr2c1*, and *Tlx* (*Nr2e1*) were enriched in the more open DARs of the 5-month-old female liver in response to As and BPA10mg exposure; among these, the expression of *Nfic* was significantly downregulated only in females under both exposures (**S-Fig.4G**). In males, CCAAT enhancer binding protein epsilon (*Cebpe*) was enriched in the more closed DARs in response to BPA10mg, PM_2.5_-CHI, and TCDD exposure. Similar sex-specific TF enrichment was also observed in the de novo motif analysis of differentially methylated regions (DMRs) (**S-Fig. 4 c-f**).

**Figure 4.**
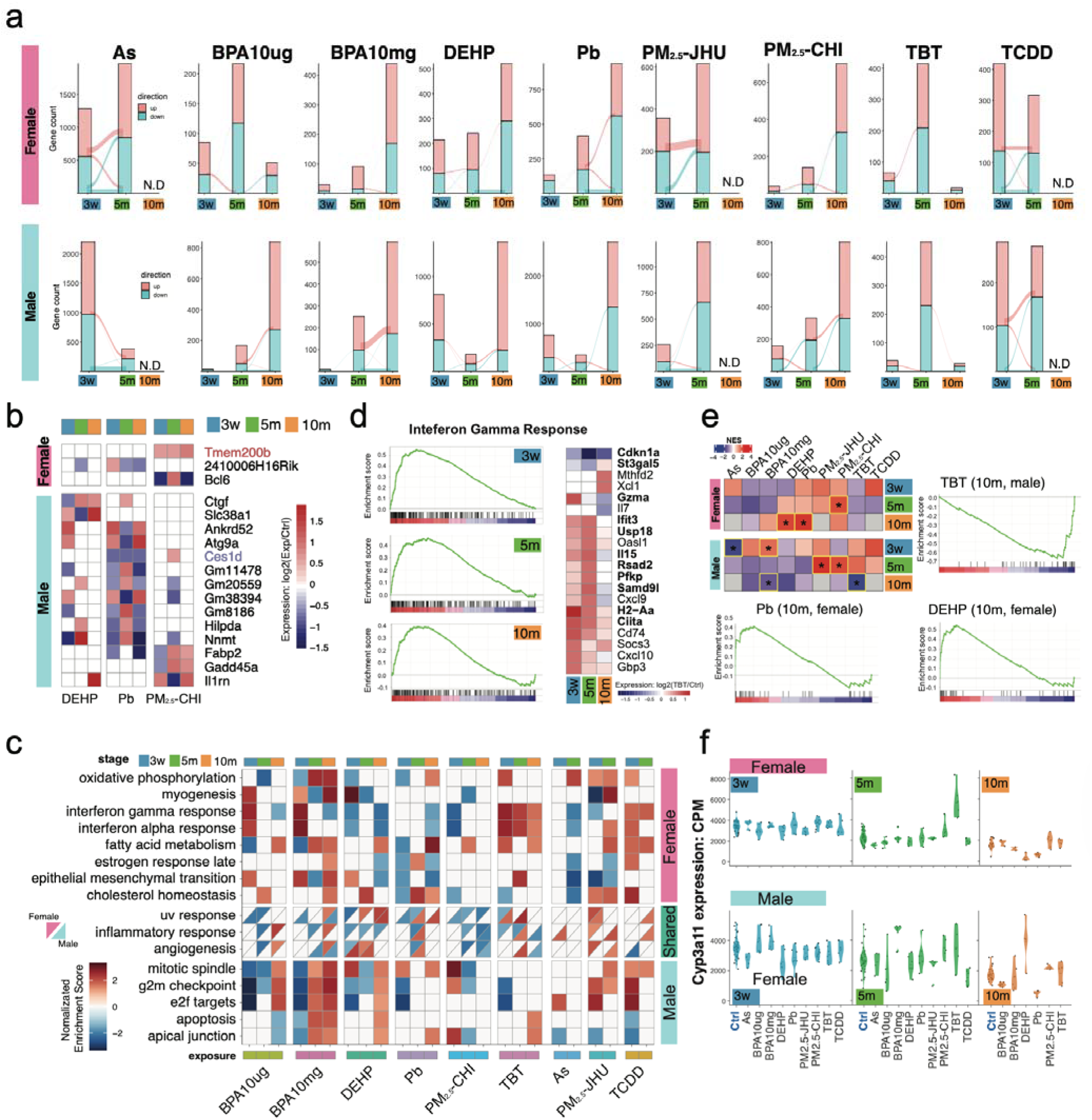
Environmental exposures induced gene expression changes across the lifespan. **a)** The number of up-regulated (pink) and down-regulated (turquoise) DEGs in female (Top) and male (bottom) in response to the same exposure at 3 weeks, 5 months, and 10 months old. The lines connecting two life stages indicated the consistent DEGs at both stages. **b)** Heatmap of expression changing (fold change) of DEGs consistently changed expression at all three stages. **c)** Significantly enriched GSEA Hallmark gene sets (FDR<0.05) in response to distinct toxicant exposures in female and male across three life stages. **d)** Enriched Hallmark gene set of Interferon Gamma response in TBT-exposed female liver at three life stages (left) and heatmap of expression changing (fold change) of DEGs in Interferon Gamma response. **e)** Heatmap of enrichment score of gene set of liver detoxification in response to distinct exposures at three stages, and the enrichment plot of TBT (male), Pb (female), and DEHP (female) at 10 month old. **f)** expression plot of *Cyp3a11* in female and male at three stages across multiple toxicant exposures.

We further explored the TFs shared between females and males under the same exposure. Interestingly, female-male shared TFs enriched in both DARs and DMRs under As exposure exhibited similar patterns: *RARa* was enriched in more closed DARs in both sexes, while *Cux2*, *CEBPE*, and *STAT5* were enriched in hypomethylated DMRs (**Fig 3d**). Similarly, TFs enriched in hypomethylated DMRs induced by PM2.5-JHU exposure were shared by both female and male. Conversely, TFs shared between females and males enriched in DARs induced by BPA10mg, TBT, and TCDD exposures exhibited reversed patterns.

### Sex-specific dynamic molecular response across the mouse lifespan

We collected transcriptomic and chromatin accessibility signatures in response to each environmental toxicant at three life stages and compared these changes across the mouse lifespan (**Fig 4a, S-Fig 5A**). In general, we observed an accumulating pattern of molecular signatures associated with the aging process. For example, during the transition from 3 weeks to 5 months in females, only TCDD exposure induced more differentially expressed genes (DEGs) at the younger stage. However, in males, As, DEHP, Pb, and TCDD exposure induced more DEGs at the younger stage, suggesting that young male animals might be more sensitive than adults. During the transition from 5 months to 10 months, TBT exposure induced fewer DEGs in both females and males. A similar reduction in DEGs was also observed in females exposed to BPA10µg. Other exposures induced more gene expression changes in the elder stage, indicating early-life exposures manifest their effects along the mouse aging process.

**Figure 5.**
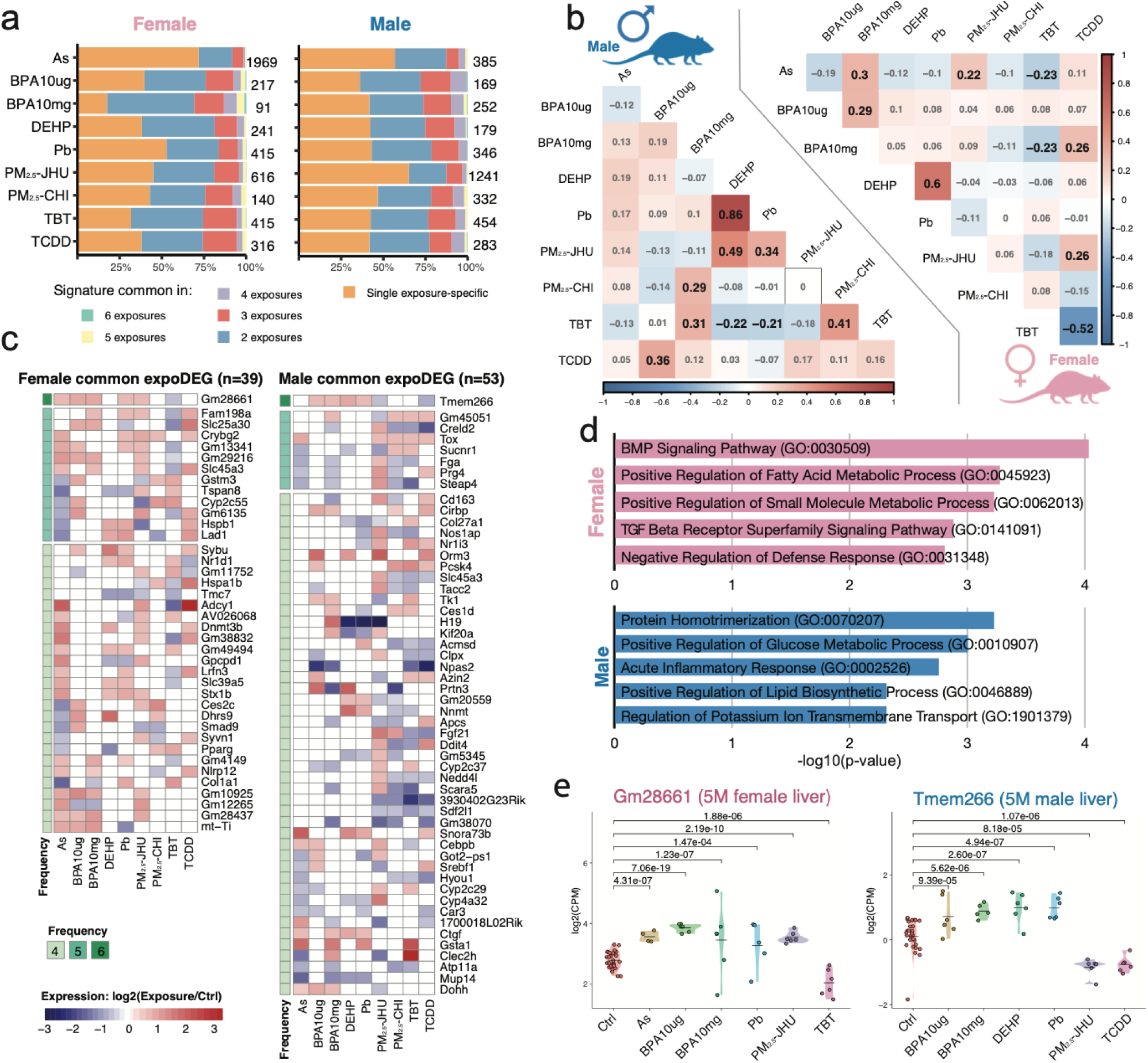
Common differentially expressed genes (DEGs) across multiple environmental toxicant exposures. **a)** Distribution of DEGs commonality in response to each toxicant exposure in 5-month-old female and male livers. The number on the right side indicated the total number of DEGs. **b)** Gene expression correlation between two exposures in male (left) and female (right). **c)** The DEGs commonly identified in multiple exposure conditions (n>3) in both sexes. **d)** Top GO biological processes enriched DEGs commonly identified in multiple exposure conditions. **e)** expression plot of *Gm28661* in female liver and *Tmem266* in male liver at 5 months old.

Surprisingly, all molecular changes, including both transcriptomic and chromatin accessibility signatures, were highly life-stage-specific, with only a very limited number of signatures maintained across different life stages. In females exposed to PM_2.5_-CHI and Pb, we identified only 2 (*Tmem200b* and *Bcl6*) and 1 (*2410006H16Rik*) differentially expressed genes (DEGs), respectively (**Fig 4b**). In males, relatively higher numbers of DEGs were found in response to DEHP, Pb, and PM_2.5_-CHI exposures: 2, 9, and 3 DEGs, respectively (**Fig 4b**). Among these genes, only two were consistently regulated across all three life stages: *Tmem200b* was consistently upregulated in PM2.5-CHI-exposed females, and *Ces1d* was consistently downregulated in Pb-exposed males (**Fig 4b**). Similarly, less consistent molecular changes with aging were also observed in the alterations of chromatin accessibility from 3 weeks to 5 months (**S-Fig. 6A**).

**Figure 6.**
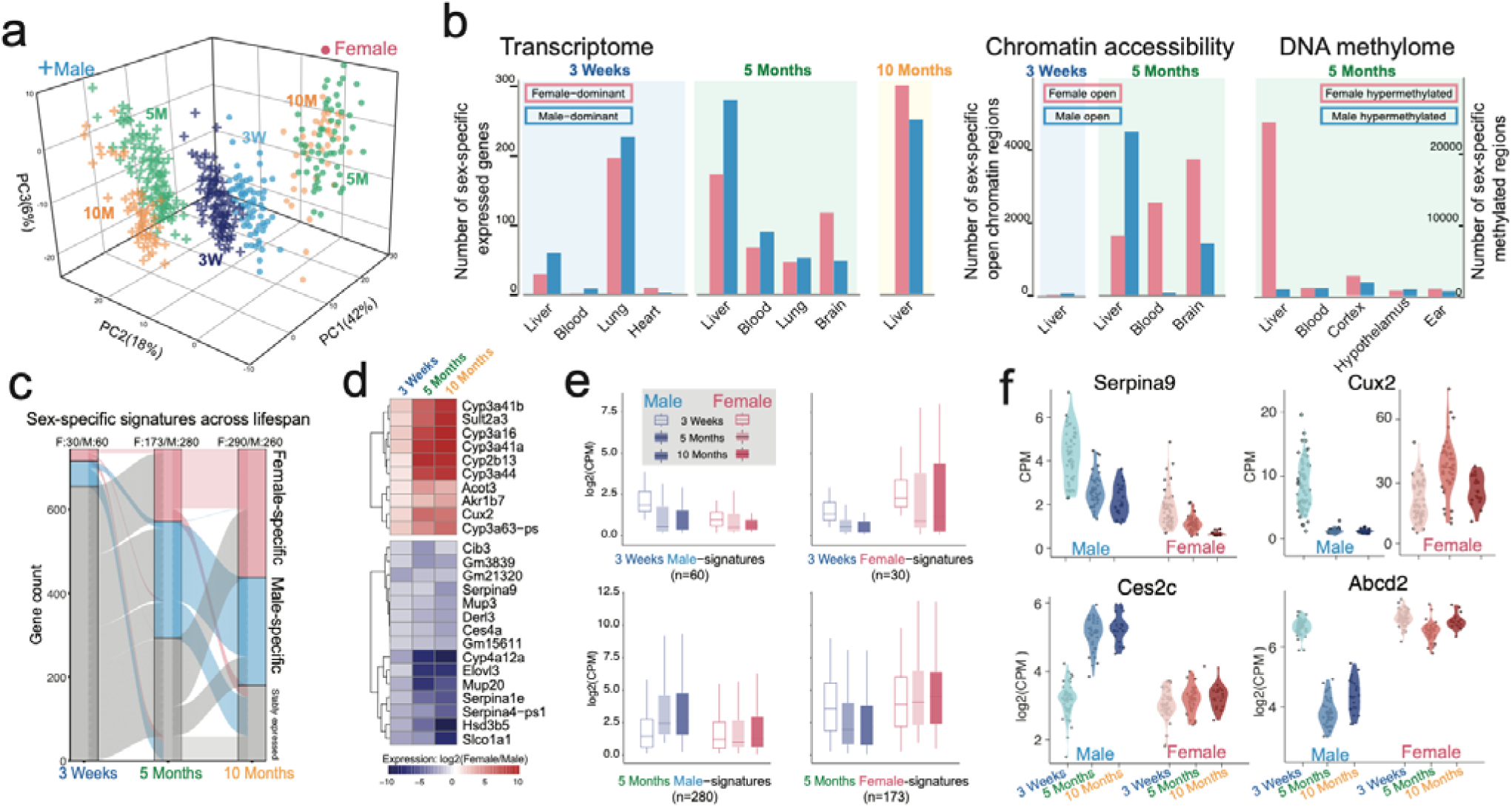
Sex-biased molecular signatures of liver at three life stages. **a)** PCA plot of liver transcriptomes of untreated female and male animals at 3 weeks, 5 months, and 10months old. **b)** Number of female-dominant (pink) and male-dominant (blue) expressed genes (left), chromatin accessible regions (middle), and differently methylated regions (right) in liver and other tissues. **c)** Dynamic changes of sex-biased expressed genes at three stages in liver. **d)** Genes consistently exhibiting female-dominant (red) or male-dominant (blue) expression patterns in mouse liver. **e)** Overall expression pattern of sex-biased expressed genes of 3 weeks and 5 months across three life stages. **f)** expression plot of *Serpina9*, *Cux2*, *Ces2c*, and *Abcd2* in female and male liver overtime.

Due to the limited continuous molecular changes (**Fig 4a**) and gene oncology enrichment (**S-Fig 5B**) observed with aging, we further investigated whether the perturbed pathways were consistent throughout aging in response to the same exposure. We performed Gene Set Enrichment Analysis (GSEA) based on transcriptomic changes in response to each exposure at three life stages and identified hallmark gene sets enriched in sex-specific and shared fashions between females and males (**Fig. 4c**). We first noticed strong sex-specific pathway enrichment patterns in the GSEA results. In females, all exposures perturbed interferon response pathways and fatty acid metabolism in distinct directions. In males, cell cycle-related pathways, including mitotic spindle and G2M checkpoints, were significantly enriched. Notably, BPA10mg exposure initially induced strong negative enrichment of these pathways at 3 weeks but dramatically stimulated positive enrichment at later life stages. We further explored the Interferon Gamma Response (IFN-γ) pathway in TBT-exposed females. Despite strong positive enrichment at all three life stages, the expression changes of the genes involved were relatively similar but not statistically significant enough to be identified as differentially expressed genes at all stages (**Fig. 4d**). Most genes belonging to the IFN-γ pathway exhibited certain inconsistent changes with aging. For example, *Cdkn1a* was downregulated at all three stages but only identified as a DEG at 5 months old.

We further examined the genes related to the liver detoxification process **(S-Fig. 5C**). The GSEA analysis of the liver detoxification gene set indicated distinct enrichment patterns, but statistical significance was observed only under limited exposure conditions (**Fig. 4e**). In females, DEHP and Pb exposure induced strong positive enrichment of detoxification genes at 10 months, and PM_2.5_-CHI induced positive enrichment at 5 months (**Fig. 4e**). In males, detoxification genes were generally negatively enriched at 10 months under all exposures, especially BPA10mg and TBT (**Fig. 4e**), suggesting that these endocrine-disrupting chemicals significantly impact males more than females. The gene *Cyp3a11*, the mouse ortholog of human *CYP3A4*, plays a crucial role in liver detoxification^60^ and shows dominant expression levels when compared to other P450 genes. We observed clear dysregulation of Cyp3a11 at all three life stages, particularly in males (**Fig. 4f**).

### Commonality of sex-specific signatures across distinct exposures

Since we observed that different exposures resulted in vastly different numbers of molecular signatures, we examined the commonality of molecular signatures (DEGs, DARs, and DMRs) across distinct exposures in both females and males. In general, both the sex-specific and shared molecular signatures were highly exposure-specific (**S-Fig-6a**). On average, in the 5-month-old liver, nearly 50% of DEGs in response to each exposure were specific to that exposure, and fewer than 25% of DEGs could be identified in at least three exposures (**Fig. 5a**). There are only 18 and 27 genes commonly changed in response to at least four exposures in female and male livers(**S-Fig-6b**). This phenomenon was similar in both male and female across all three life stages (**S-Fig. 7-8**). We further examined the correlation of transcriptome changes across distinct exposures for each sex. Based on the expression correlation of the union of DEGs in all exposure conditions, we noticed distinct correlations between different exposures (**Fig 5b**). For example, DEHP and Pb exposure induced strong positive correlation of transcriptomic changes which were positively correlated in males, but correlation was relatively weak in females, suggesting both exposures might significantly impact males in a similar pattern rather than females. In male, we observed that PM_2.5_-JHU exposure strongly correlated with DEHP and Pb, while TBT exposure positively correlated with BPA10mg exposure but negatively correlated with DEHP. In females, TBT exposure was strongly negatively correlated with TCDD exposure, suggesting these two exposures induced oppositely expressed changes in females.

**Figure 7.**
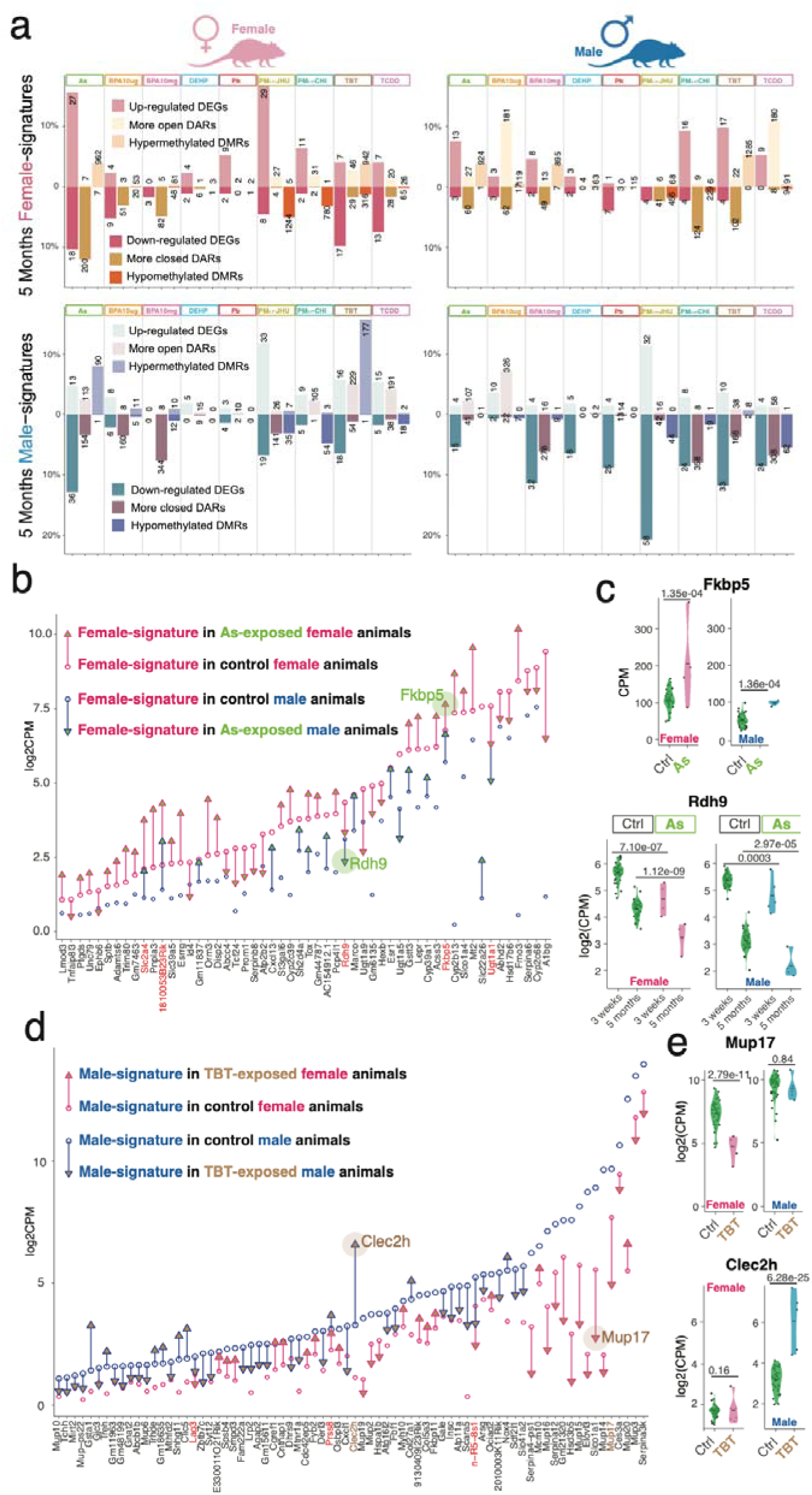
Early-life exposures to toxicants altered the sex-biased expression patterns. **a)** Numbers of DEGs, DARs, and DMRs belong to female-dominant expressed genes (top) and male-dominant expressed genes (bottom) of 5-months-old liver in female (left) and male (right) under distinct exposures. **b)** Significant expression changes of female-dominant expressed genes in female (pink) and male (blue) animals exposed to early-life As exposure. **c)** expression plot of *Fkbp5* and *Rdh9* in control and As-exposed animals. **d)** Significant expression changes of male-dominant expressed genes in female (pink) and male (blue) animals exposed to early-life TBT exposure. **e)** expression plot of *Mup17* and *Clec2h* in control and As-exposed animals.

Such anticorrelated regulation was also observed in the commonly identified DEGs. We focused on the 39 and 53 genes that were dysregulated in at least four exposures in females and males, respectively (**Fig. 5c**). In both sexes, we noticed that the genes commonly altered by DEHP and Pb exhibited generally consistent changes. For instance, in females, genes such as *Lad1*, *Sybu*, and *Nr1d1* were consistently dysregulated, and in males, the genes *H19*, *Kif20a*, and *Ctgf* showed similar patterns of dysregulation. In females, genes commonly changed by TBT and TCDD were always oppositely regulated, including *Fam198a* and *Adcy1*. In both sexes, the gene signatures dysregulated by the two PM_2.5_ exposures were poorly correlated, suggesting that the liver responds uniquely to distinct components of PM_2.5_ (**Fig.5a, S-Fig.6d**). In males, PM2.5-JHU, which contained a high amount of sulfite (S), induced a similar molecular response to DEHP and Pb exposure, while PM_2.5_-CHI, which contained high levels of iron (Fe) and sodium (Na) components, induced a molecular response similar to TBT exposure. Gene ontology enrichment analysis suggested that the 39 genes commonly dysregulated in at least four exposures in females were associated with BMP/TGF-beta signaling and metabolic processes. In males, the 53 commonly dysregulated genes were related to protein homotrimerization, glucose metabolism, and inflammatory response (**Fig. 5d**). We identified a single gene for each sex that was dysregulated in six exposures. In females, a pseudogene associated with the mitochondrial oxidative phosphorylation process^61^ was upregulated in As, BPA10mg, Pb, and PM_2.5_-JHU exposures but downregulated by TBT exposure. In males, a zinc-activated voltage sensor^62^ was upregulated by BPA, DEHP, and Pb exposures, but downregulated by PM2.5-JHU and TCDD exposures (**Fig. 5e**).

### Sex-biased expressed molecular signatures of the liver across lifespan

During normal liver development, sex-biased patterns are highly associated with developmental stages and shape liver-specific functions, including metabolism, immune response, and aging. To understand how the sex-biased transcriptome develops across different life stages, we first integrated the transcriptomes of three stages in both female and male livers in PCA space. We observed a clear but minor separation between the sexes already at 3 weeks old (**Fig 6a, S-Fig-9a**). By 5 months old, the sex-biased expression pattern became well-established and remained consistent at 10 months old. We identified sex-biased genes in the liver at all three stages and observed that the number of DEGs increased dramatically at 5 months and stabilized at 10 months, while different trends were observed in other tissues (**Fig. 6b, S-Fig-9b,c**). For example, there were more than 400 sex-biased genes in the lung tissue at 3 weeks old, but only 100 DEGs at 5 months old. Compared to other tissues, the liver exhibited the strongest sex-biased expression pattern at 5 months of age. Similarly, we identified approximately 6,000 DARs between female and male livers at 5 months old, with very few chromatin accessibility differences at 3 weeks old (**S-Fig-10a**). Notably, we found more open DARs in females in blood and brain tissues, whereas more open DARs were identified in male liver tissue. At the DNA methylation level, the female liver exhibited a strong hypermethylation pattern compared to males, and other tissues did not show a strong sex-biased methylation pattern (**Fig. 6b, S-Table 17-19**).

The comparison of sex-biased expressed liver genes at three life stages revealed unexpected patterns. Most genes significantly differentially expressed at 3 weeks were not sex-biased after maturation, while the majority of sex-biased expressed genes at 5 months were established and remained during adulthood (**Fig. 6c**). We identified only 10 genes in females and 15 genes in males that maintained sex-biased expression patterns across all three life stages, including many P450 genes (**Fig. 6d**). Further exploration of these stage-specific sex signatures indicated intriguing changing patterns. We noticed that male sex signatures changed more dynamically compared to female signatures (**Fig. 6e, S-Fig-10b.**). In males, genes such as *Serpina9* were highly expressed at 3 weeks but downregulated at later stages, while these genes were generally lowly expressed in female livers across all three stages (**Fig. 6f**). In contrast, female signatures, such as *Cux2*^63^, were highly expressed at 3 weeks and many of them continued to remain highly expressed in female livers at later stages, whereas these genes were generally lowly expressed in male livers (**Fig. 6f**). The majority of 5-month sex signatures indicated similar expression levels at 3 weeks for both females and males (**Fig. 6e**). However, male signatures, such as *Ces2c*^64^, were significantly upregulated at 5 and 10 months in males but maintained 3-week expression levels in females at later stages. Conversely, female signatures, such as *Abcd2*^65^, maintained 3-week expression levels in females at later stages but were significantly downregulated in males at 5 and 10 months. Together, this data suggests that the sex-biased expression pattern in the liver is mainly due to dynamic expression changes in males, while the temporal expression pattern in females is more stable.

### The impacts of environmental toxicants on the expression of liver sex signatures

We examined the dysregulation of sex signatures in response to early-life exposure to environmental toxicants in 5-month-old mice and observed distinct impacts in an exposure-specific manner (**Fig. 7a**, **S****-Fig-10c, S-Table 20-22**). In females, early-life arsenic (As) exposure induced changes in 45 female-biased genes at 5 months, resulting in 27 upregulated and 18 downregulated genes (**Fig. 7b**). Additionally, 200 female-open DARs lost chromatin accessibility and became more closed under exposure, along with an increase in percentage hypermethylation at 962 female-biased DMRs. Some of the female highly expressed genes were also dysregulated in exposed male livers. For example, *Fkbp5*, a protein involved in regulating the stress response and associated with alcohol-associated liver disease^66^, was significantly upregulated in both female and male livers in response to As exposure (**Fig. 7c, d**). *Rdh9*, a key component of the retinoid metabolic pathway^67^, was highly expressed in females and significantly downregulated in both sexes in response to As exposure (**Fig. 7c, d**).

In males, we observed that As, BPA10mg, PM_2.5_-CHI, TBT, and TCDD exposure induced more upregulation of female-biased signatures, which were generally lowly expressed in normal male livers. Moreover, As, BPA10mg, DEHP, Pb, PM_2.5_, TBT, and TCDD exposure induced more downregulation of male-biased signatures, which were highly expressed in normal males (**Fig. 7a**, **S****-Fig-10c**). Combining these results, early-life toxicant exposure seems to impact males in a relatively consistent fashion: downregulating male signatures while upregulating female signatures. In females, we did not observe as clear a pattern. For example, Pb exposure induced downregulation of 25 and 4 male signatures in male and female livers, respectively, along with upregulation of 4 and 3 male signatures in males and females, respectively (**Fig. 7a**). TBT exposure induced downregulation of 38 and 18 male signatures in male and female livers, respectively, along with upregulation of 10 and 18 male signatures in males and females, respectively (**Fig. 7a, d**). Particularly in females, most of the relatively highly expressed genes were downregulated, such as *Mup17*, a member of the lipocalin family involved in lipid transport^68^, which was significantly downregulated in females (**Fig 7e**). In males, with many downregulated male signatures, some genes, such as *Clec2h*, associated with liver inflammation after injury^69^, were significantly upregulated (**Fig 7e**), suggesting potential liver damage induced by TBT exposure.

## Discussion

Liver morphogenesis in mice is a sexually dimorphic process involving the establishment of sex-specific molecular signatures and functional capacities during the organ’s maturation. This process begins postnatally and is primarily driven by hormonal signaling, particularly androgens and estrogens, as well as growth hormone signaling, which diverges significantly between males and females starting at puberty. As such, early-life exposure to environmental toxicants can profoundly disrupt liver morphogenesis by dysregulating the transcriptome and epigenome during this sensitive developmental window.

In this study, we leveraged a comprehensive multi-omics dataset generated by the TaRGET II Consortium to systematically examine sex-specific molecular responses, including altered gene expression, chromatin accessibility, and DNA methylation patterns, in male and female mice across three life stages. Our findings reveal that toxicant exposure from prenatal stages through postnatal week 3 can induce significant and lasting molecular alterations, including persistent transcriptional dysregulation and epigenetic remodeling. These significant long-term changes were observed in the 5-month and 10-month-old mice, corresponding approximately to 30-year-old and 50-year-old humans, respectively. Together, these results highlight the critical vulnerability of early developmental stages to environmental exposures and emphasize the need for preventive strategies to minimize toxicant exposure during early life, thereby protecting long-term liver health and maintaining normal sex-specific molecular programming.

Although most molecular changes induced by environmental toxicants are sex-specific, we identified a subset of shared molecular signatures between females and males in response to the same toxicant exposure. These shared features included both transcriptomic and epigenetic alterations. Notably, we observed more consistent patterns in DNA methylation changes across sexes, while chromatin accessibility changes generally exhibited a negative correlation with DNA methylation, supporting the well-established model of opposing roles between these two epigenetic regulatory mechanisms. Interestingly, As-exposure induced a particularly strong set of female-male-shared signatures, with positively correlated changes observed at the gene expression, chromatin accessibility, and DNA methylation levels, highlighting the persistent and widespread impact of early-life As-exposure on both transcriptional regulation and epigenetic remodeling, and suggesting the potentially harmful and long-lasting effects of arsenic, even at early developmental stages.

We also compared the disrupted KEGG pathways, biological processes, and upstream TF regulators in response to the same toxicant exposure in both female and male mice. Consistent with the observations at the level of individual molecular features, we found pronounced sex-specific patterns in these higher-order regulatory mechanisms. While certain pathways, such as hepatocellular carcinoma, immune response, estrogen signaling, cell cycle regulation, and focal adhesion were commonly perturbed in both sexes across various life stages, a closer examination of the gene-level changes within these pathways revealed striking sex-specific expression patterns. These findings indicate that, despite some shared pathway-level disruptions, the underlying regulatory responses are fundamentally sex-dependent, indicating the sex-specific regulation as a dominant feature in the molecular response to environmental toxicant exposure.

Further analysis revealed that only a very small number of exposure-induced molecular signatures were consistently altered throughout mouse development and aging, indicating that the majority of toxicant-induced changes were highly stage-specific. In most of the exposures, molecular disruptions observed at early life stages were not present in adult animals. This suggests that mice possess a robust recovery capacity, and the liver can partially restore normal gene expression patterns over time following early-life exposure to toxicants. However, despite this apparent recovery, early-life exposure still resulted in stronger long-term molecular and functional consequences, such as disrupted liver metabolism^70^. In most of the exposures, the effects of toxicant-induced dysregulation persisted or re-emerged later in life, particularly at 10 months of age, indicating cumulative stress or damage. These findings suggest a dual dynamic: while some genes and pathways may normalize, more may exhibit latent dysregulation, ultimately contributing to chronic liver dysfunction, increased disease risk, or age-related pathologies.

Finally, we examined sex-biased molecular patterns in untreated (control) animals to assess the natural trajectory of liver sex morphogenesis. As expected, we observed clear sex-specific gene expression as early as 3 weeks of age, which became more fully established by adulthood (5 months). Interestingly, these sex-biased patterns also showed stage-specific dynamics. Many genes that were sex-biased at 3 weeks lost their sex-specificity at later time points, indicating that early sex differences are not always maintained into adulthood. Consistent with previous finding^71^, at both 3 and 5 months, we found that sex-biased expression was primarily driven by dynamic changes in male livers, whereas female livers tended to maintain more stable expression patterns over time. This suggests that the male liver may have higher transcriptional plasticity, potentially making it more sensitive and responsive to environmental perturbations, which was supported by our observation that male animals in general exhibited a greater number of exposure-induced molecular alterations compared to females. Moreover, in males, early-life toxicant exposures disrupted normal sex-biased gene expression, often leading to a loss of male-specific gene signatures and, in some cases, the acquisition of female-like expression profiles. In contrast, female animals showed less consistent or pronounced alterations, with both male-specific and female-specific signatures being partially disrupted. As male-biased hepatic gene expression is governed in large part by pulsatile growth hormone secretion, future work should examine effects of developmental toxicant exposures on this pathway^71^. These findings suggest that early-life environmental exposures can compromise the establishment and maintenance of sex-specific liver identity, particularly in males, which may contribute to sex-specific disease vulnerabilities later in life.

## Methods

### Animals and Exposures

C57BL/6 (B6; Jackson Laboratory, Bar Harbor, ME) mice were used for all experiments, except for the lead (Pb) and phthalate (DEHP) exposure studies, which employed wild-type non-agouti (a/a) mice derived from a >230-generation colony of viable yellow agouti (Avy) mice.

Mice were exposed to toxicants perinatally via maternal diet, drinking water, or air breathing, with the exposure window spanning from pre-conception through weaning. In brief, two weeks before mating, virgin female dams (6–8 weeks old) were randomly assigned to the exposure or control group. Detailed methods for the animal exposure and tissue collection were described in our previous study^72^.

### Raw sequence data and processing

In total, 493 RNA-seq, 346 ATAC-seq, and 186 WGBS data from liver samples of weaning (3 weeks), early adulthood (5 months), and late adulthood (10months) (Supplementary table 1) were gathered from mice exposed to pollutants during early development, including arsenic (As), lead (Pb), bisphenol A (BPA), tributyltin (TBT), di-2-ethylhexyl phthalate (DEHP), 2,3,7,8-tetrachlorodibenzo-p-dioxin (TCDD), and particulate matter less than 2.5 µm in diameter (PM2.5), and their age-matched non-expsure controls. All the raw-data were provided at TaRGET II data portal (https://data.targetepigenomics.org/).

Raw fastq files of RNA-seq data were processed by Cutadapt^73^ (v1.16; --quality-cutoff=15,10 -- minimum-length=36), FastQC^74^ (v0.11.7), and STAR^75^ (v2.5.4b; --quantMode TranscriptomeSAM -- outWigType bedGraph --outWigNorm RPM) to do the adapter trimming, generating QC report and mouse genome mapping (mm10) using our own built pipeline, the TaRGET-II-RNA-seq-pipeline (https://github.com/Zhang-lab/TaRGET-II-RNAseq-pipeline). Next, featureCounts (v1.5.1)^76^ was used to calculate the gene expressions across normal and exposure samples within the pipeline based on GENCODE vM20 gene annotation of mouse genome^77^.

ATAC-seq fastq files were processed by our own built pipeline based on AIAP^78^, the TaRGET-II-ATAC-seq-pipeline (https://github.com/Zhang-lab/TaRGET-II-ATACseq-pipeline) that integrated AIAP packages, including optimized QC reports and analysis pipeline with default parameters to generate the open chromatin regions (OCRs). Then, the consensus regions of OCRs across all exposure and control samples were generated with Index (https://github.com/Altius/Index) method used for the downstream analysis^79^. The ATAC-seq signals of consensus OCRs were calculated by using the intersectBed method of bedtools^80^.

The TaRGET-II-WGBS-pipeline (https://github.com/Zhang-lab/WGBS_analysis) was built by using the Cutadapt^73^ (v1.16; --quality-cutoff=15,10 --minimum-length=36) and Bismark^81^ (v0.19.0; --bowtie2 -X 1000 --score_min L,0,-0.6 -N 0 --multicore 2 -p 4) to do the trimming and mouse genome mapping. The pipeline also incorporated quality control, generated user-friendly files for computational analysis, and output genome browser tracks for data visualization.

The RNA-seq and ATAC-seq datasets were normalized to correct for batch effects arising from different data production centers by using the method described in the flagship paper, and then the consortium-normalized read count data were used for downstream analysis.

### Differential Transcription and Epigenomic Analysis

The differentially expressed genes (DEGs) between control and exposure samples under different conditions were identified by using DESeq2^82^ (v1.34.0) as previous studies^83,84^. Consortium-normalized read count data were used to identify the DEGs between the exposed and control samples . Genes were considered significantly differentially expressed if they met the following stringent criteria: an adjusted p-value (Benjamini-Hochberg correction) less than 0.001 and an absolute log2 fold change greater than log2(1.5), equivalent to a 1.5-fold change in expression.

Differentially accessible regions (DARs) between control and exposure samples were identified using edgeR (v3.36.0)^85^, as previous study^86,87^. Consortium-normalized ATAC-seq reads were used to identify DARs, with significance determined by the following cutoffs: an absolute log2 fold change greater than log2(1.5) (a 1.5-fold change in accessibility) and a false discovery rate (FDR) less than 0.01.

Differential methylation regions (DMRs) between control and exposure samples were identified using a method adapted from previous studies^88,89^. Methylation levels were quantified using whole-genome bisulfite sequencing (WGBS) data. Briefly, raw WGBS reads were aligned to the reference genome using Bismark (v0.23.1) with default parameters, followed by deduplication to remove PCR artifacts.

Methylation levels at individual CpG sites were calculated as the ratio of methylated reads to total reads (methylated + unmethylated) at each site, with a minimum coverage of 10 reads per site to ensure reliability. DMRs were identified through a genome sliding approach, as previous studies^89,90^: the genome was divided into sliding 200bp windows, the methylated and unmethylated reads counts in each window were calculated in both control and exposed samples, and a Chi-squared test was applied to identify DMRs. Regions were defined as differentially methylated if they exhibited a methylation level difference of at least 0.1 (a 10% absolute difference in methylation between control and exposed condition) and a Q-value (adjusted for multiple testing using the BH method) less than 0.1. Additionally, only regions with at least 2 CpG sites and a minimum average coverage of 10 reads per site were considered to enhance statistical power and reduce noise in the analysis, and the neighboring regions were merged to maximize the width of DMRs.

### Gene Ontology (GO) Enrichment Analysis

To investigate the biological processes affected by exposure-induced differentially expressed genes (DEGs), we performed GO enrichment analysis using the enrichGO function from the clusterProfiler R package. Gene symbols of DEGs were used as input, with the following parameters: ont = “BP” (Biological Process), pAdjustMethod = “BH” (Benjamini-Hochberg correction), pvalueCutoff = 0.01, and qvalueCutoff = 0.05. Only GO terms containing at least five DEGs were retained for downstream analyses to ensure statistical robustness and biological relevance. This analysis enabled the identification of significantly enriched biological pathways and functional categories associated with each toxicant exposure in both female and male liver samples across developmental stages.

### KEGG Pathway Enrichment Analysis

To identify biological pathways significantly affected by toxicant exposure, we performed Kyoto Encyclopedia of Genes and Genomes (KEGG) enrichment analysis using the run_pathfindR function from the pathfindR R package. For each exposure condition, a combined list of upregulated and downregulated DEGs was used as input. The input data included gene symbols, corresponding log_₂_ fold changes, and adjusted p-values derived from differential expression analysis. The analysis was performed with the following parameters: a minimum gene count of 5 per pathway, and a p-value threshold of 0.1 to retain enriched pathways. This approach enabled the identification of exposure-associated signaling pathways and helped characterize both shared and sex-specific regulatory responses across different toxicants and life stages.

### Gene Set Enrichment Analysis (GSEA)

To evaluate the coordinated transcriptional responses to early-life environmental toxicant exposures, Gene Set Enrichment Analysis (GSEA) was performed using GSEABase (v3.1) in r environment. For each exposure condition, genes were ranked based on log_₂_fold change values derived from differential expression analysis using DESeq2. Genes with low expression (mean CPM < 1) and located on sex chromosomes (chrX and chrY) were removed across all samples and excluded from ranking. Curated gene sets were obtained from the Molecular Signatures Database (MSigDB v7.5), and MH: hallmark gene sets were used. Hallmark gene sets summarize and represent specific well-defined biological states or processes and display coherent expression. For GSEA enrichment calculation, parameters were set as follows: nPerm = 1000; minGSSize = 15; maxGSSize = 500; pvalueCutoff = 0.05; pAdjustMethod = “BH” (Benjamini–Hochberg correction). Significantly enriched gene sets (FDR < 0.05) were visualized using enrichment plots, dot plots, and ridge plots to highlight coordinated upregulation or downregulation of biological processes. Enrichment scores, normalized enrichment scores (NES), and leading-edge subsets were used to interpret key drivers of pathway-level responses.

To assess the coordinated expression changes of liver detoxification-related genes in response to environmental toxicant exposure, we performed Gene Set Enrichment Analysis (GSEA) focusing on cytochrome P450 (CYP) genes. A curated list of 55 CYP genes involved in detoxification pathways was obtained from the study “Detoxification Cytochrome P450s (CYPs) in Families 1–3 Produce Functional Oxylipins from Polyunsaturated Fatty Acids” and used as a reference gene set. For the GSEA, genes were ranked using the Signal-to-Noise (S2N) metric based on their expression profiles across exposure and control groups. The analysis excluded gene sets with fewer than 15 or more than 500 genes to improve statistical reliability. The number of permutations was set to 100 to assess significance. For visualization, the top 20 enriched gene sets for each phenotype (i.e., upregulated or downregulated in exposed vs. control) were selected for plotting enrichment curves and leading-edge analyses.

### The changes of molecular features in the liver under different exposures

The ggplot2 package of R was used to generate bar plots to show number of DEGs, DARs and DMRs across exposures in liver ^91,92^. And the enriched biology processes in different exposures based on DEGs were generated by clusterProfiler package of R using the mouse annotation database (org.Mm.eg.db)^93^. The heatmap function and UpSetR package of R was used to visualize the distribution of DEGs identified in multiple exposures^94^. The intersectBed method of bedtools was used to identify the overlapped DMRs (at least 1 bp) between different exposures, or between female and male, as the common signatures. Next, the coefficient correlation between females and males in different exposures was separately calculated by using the cor function of R based on changes of DEGs, DARs, and DMRs under exposures in females and males. The transcription factors (TFs) and epigenetic modification factors (EpiGenes) were separately gathered from the AnimalTFDB3.0 and Epi-Modifiers databases^95,96^. The differentially expressed TFs (DE-TF) and EpiGenes (DE-EpiGenes) were identified and assigned into different types based on the two downloaded annotation files.

### The common features between female and male in response to different exposures

To investigate sex-independent molecular responses to environmental toxicant exposures, we identified common features between female and male animals for each exposure separately. Differentially expressed genes (DEGs) that were significantly altered in both sexes under the same exposure condition were defined as female–male-shared DEGs, based on consistent gene symbols annotated by GENCODE vM20. Shared chromatin accessibility changes (DARs) were identified as female–male-shared DARs if the genomic coordinates of differential peaks were identical between sexes. For differentially methylated regions (DMRs), shared elements were defined as regions with at least 1 bp of overlap between female and male DMRs using the intersectBed function from BEDTools. All shared molecular features (DEGs, DARs, DMRs) were processed and compared using custom R scripts, and their distribution across exposures was visualized with ggplot2. To explore the functional relevance of the shared DEGs, we conducted biological process enrichment analysis using the Enrichr web-based tool. Gene Ontology (GO) terms for biological processes were analyzed using the Enrichr API^97^, and significantly enriched terms (adjusted p-value < 0.05) were retained for downstream interpretation.

### *De novo* transcription factors binding motifs analysis

The transcription factor (TF) binding motifs enriched in differentially accessible regions (DARs) — both those with increased and decreased accessibility — as well as in hypermethylated and hypomethylated differentially methylated regions (DMRs) following exposures, were analyzed separately for female and male samples. This analysis was conducted using the findMotifsGenome.pl script with the -size given option from HOMER software (v4.11.1) ^98^. Significantly enriched de novo binding motifs were identified based on stringent criteria calculated by HOMER: The transcription factor binding site (TFBS) must be present in at least 5% of the accessible DARs within the exposure group. The match score for the TFBS motif must exceed 0.8, indicating a high confidence in motif similarity. The associated P-value must be less than 1 × 10^-11^, ensuring statistical significance. Following motif identification, known transcription factor genes (TFs) predicted to bind these enriched TFBS were extracted, requiring a motif match score greater than 0.8 to ensure accuracy. These TFs were subsequently classified using annotation files obtained from the AnimalTFDB 3.0 database^99^, enabling categorization into TF families and functional classes.

To explore tissue specificity and overlap, the number of DARs enriched for common TFBS between liver and blood samples was quantified and visualized using the jvenn tool, facilitating the identification of shared regulatory elements across tissues. Additionally, sequence logos representing the TFBS motifs were retrieved from the JASPAR database^100^ to visually summarize the nucleotide composition and conservation patterns of the identified binding sites. For functional interpretation, DARs containing these TF binding motifs were isolated from the HOMER results. The genes proximal to these DARs in different exposure conditions were then subjected to Gene Ontology (GO) enrichment analysis using the enrichGO function from the clusterProfiler R package. This allowed identification of biological processes significantly associated with the TF-regulated regions. Finally, enriched GO terms were visualized using the emapplot function of clusterProfiler, providing an interpretable network representation of related biological processes impacted by the exposures.

### GSEA of liver detoxication genes

Genes of CYPs in detoxification pathways were retrieved from a previous work [Detoxification Cytochrome P450s (CYPs) in Families 1–3 Produce Functional Oxylipins from Polyunsaturated Fatty Acids] (n=55), and used as reference in GSEA analysis. For GSEA setting, Metric for ranking genes was Signal2noise, sets with size of more than 500 or fewer than 15 were excluded, the number of markers as 100, and top 20 sets of each phenotype were used to plot graphs.

### WGBS QC

We excluded CpG sites with <10x coverage. Samples with <15M qualified CpG sites are not included in following analyses. Samples with proportion of CpG sites that have methylation levels >0.7 less than 60% are excluded. We calculate sex identification index (read counts of chrY/sum of reads counts of chrX or chrY) for each sample. We excluded samples that have the index not match to their sex labels. A total of 186 samples remained after filtering.

### DMR analysis

WGBS data were processed with the pipeline. The single C were processed with surrounding C on the other strand to get higher coverage of CpG sites. Only CpG sites with ≥10x coverage are used to calculate the number of methylated and unmethylated reads of each 200-bp window along the genome.

Chi-square test was performed with methylated and unmethylated reads of exposure versus control conditions or female control versus male control. In each window, CpG sites found in all replicates of an exposure and in 80% of all control replicates under the same condition are used in chi-square test. The windows with methylation level difference greater than 0.1 and q<0.1 in chi-square test were used to merge with adjacent windows if they are both hypermethylated or hypomethylated. Regions with ≥2 CpG sites satisfying these cutoffs are considered as DMRs.

### Correlation study

For each sex, matrix of log2-fold change of exposure DEGs that can be found in≥2 exposures are used to perform correlation test with “cor” function. For exposure DEGs that are not significant under an exposure, the log2-fold change values are used instead. The correlation matrix was generated using colorRampPalette function.

### GO enrichment analysis

Gene symbols of exposure DEGs were used as input to perform biological processes analysis with “enrichGO” function in “clusterProfiler” package. The setting is “ont=“BP”, pAdjustMethod = “BH”, pvalueCutoff = 0.01, qvalueCutoff=0.05”, only GO terms with >=5 genes were used in the following analyses.

### KEGG analysis

Information of gene symbols, log2-fold change and adjusted p-values of Up-regulated and down-regulated DEGs by exposure together were used as input in KEGG analysis with “run_pathfindR” function in “pathfindR” package. The minimum gene number is each KEGG pathway was 5, and lowest p-value cutoff is 0.1.

### Enrichment score calculation

Enrichment score is determined by fraction of number of sex-specific DEGs in an exposure among sex-specific DEGs divided by fraction of exposure DEGs among total expressed genes. Hypergeometric test was used to determine the probability of obtaining at least a specific number of sex-specific DEGs in exposure DEGs when sampling from total expressed genes without replacement. The p-value<0.001 was considered as significance.

## Data availability

RNA-seq, ATAC-seq, and WGBS data in the paper have been deposited through Gene Expression Omnibus (GEO) repository GSE146508, and those data are also available at the TaRGET II data portal (https://data.targetepigenomics.org/). All analysis results are visualized at the accompanying database ToxiTaRGET https://toxitarget.com/. All the code used in this study is deposited to GitHub (https://github.com/Zhang-lab/TaRGET-II-sex-specific-molecular-response-to-early-life-exposures-of-toxic-substances/)

## Acknowledgements

This work was supported by the NIEHS as part of the Toxicant Exposures and Responses by Genomic and Epigenomic Regulators of Transcription II (TaRGET II) Consortium through U24ES026699 (T.W.), U01ES026697 (D.D), U01ES026719 (C.L.W., M.S.B., and T.W.), U01ES02672 (S.B., S.R., and W.T.), U01ES026718 (G.M.), and U01ES026717(D.A.). This work was also supported by NIH through R35GM142917 (B.Z.), P30ES017885 and R35ES031686 (L.S., D.D.), R01 S028802 (J.C.), P30ES030285 (C.L.W.), U24HG012070 (T.W.), U41HG010972 (T.W.), and U24NS132103 (T.W.), K01 ES032048 (L.S.). This work was also supported by Chan Zuckerberg Initiative (B.Z.), Diana Helis Henry Medical Research Foundation (69760-I, C.L.W.), and Michigan Biological Research Initiative on Sex Differences in Cardiovascular Disease (M-BRISC, L.S.).

We acknowledge program leadership by members of the NIEHS TaRGET II workgroups, especially Fred. L. Tyson, Kim McAllister, Christopher G. Duncan, Amanda Garton, Lisa H. Chadwick, Maya Evanitsky.

## TaRGET II consortium (alphabetical)

### Integrative analysis leads and coordination

Benpeng Miao, Ting Wang, Bo A. Zhang,

### Integrative analysis

Cristian Coarfa, Justin A. Colacino, Shuhua Fu, Ravindra Kumar, Prashant Kumar Kuntala, Bongsoo Park, Wanqing Shao, Laurie K. Svoboda

### Integrative data production and processing

Gregory E. Crawford, Robert B. Hamanaka, Claudia Lalancette, Daofeng Li, Shaopeng Liu, Benpeng Miao, Heather B. Patisaul, Maureen A. Sartor, Tim Wiltshire, Xiaoyun Xing, Bo A. Zhang

### TaRGET II Data Production and Processing contributor

Nicole E. Allard, Raymond G. Cavalcante, Rengul Cetin-Atalay, Yujie Chen, Youngshim Choi, Alan Du, Elisa Ruiz-Echartea, Jackson P. Fredenburg, Tianyi Fu, Jaclyn M. Goodrich, Sandra L. Grimm, Silas Hsu, Brian M. Horman, Yiran Hou, Rahul Jangid, Yan Jin, Tamara R. Jones, Tiffany A. Katz, SunHong Kim, Prashant Kumar Kuntala, Yemin Lan, Bethany Latham, Tandao Li, Yan Li, Sydney Lierz, Siyu Liu, Juheon Maeng, Angelo Y. Meliton, Rachel K. Morgan, Kari Neier, Jackson P. Parker, Bambarendage P.U. Perera, Jayant M. Pinto, Deepak Purushotham, Dhivyaa Rajasundaram, Palanivel Rengasamy, Christine A. Rygiel, Alexias Safi, Erica Pehrsson, Kaitlyn A. Sun, Vinesh Vinayachandran, Kai Wang, Parker S. Woods, Bonnie HY Yeung, Jinhu Yin, Yu Zhang, Xiaoyu Zhuo

### Co-principal investigators

Cristian Coarfa, Gregory E. Crawford, Heather Lawson, Michael Province

### Scientific program management

Maya Evanitsky, Kimberly A. McAllister, Alice Zorn, Frederick L. Tyson

### Principal Investigators

David Aylor, Marisa S. Bartolomei, Shyam Biswal, Dana C. Dolinoy, Gökhan M. Mutlu, Sanjay Rajagopalan, Wan-Yee Tang, Cheryl Lyn Walker, Ting Wang

## Supplementary files

